# Role of adipocyte angiotensinogen or angiotensin type 1a receptors in the development of diet-induced atherosclerosis or angiotensin II-induced abdominal aortic aneurysms

**DOI:** 10.1101/2025.05.05.652275

**Authors:** Yasir AlSiraj, Kelly Putnam, Victoria L. English, Charles M. Ensor, Lisa A. Cassis

**Author notes:** These authors contributed equally to this research (Yasir AlSiraj, Kelly Putnam). **Corresponding author:** Lisa A. Cassis, PhD Professor, Department of Pharmacology and Nutritional Sciences College of Medicine, University of Kentucky 900 S. Limestone, Lexington, KY 40536-0200. **Author contributions:** YA (data curation, formal analysis, writing – review and editing), KP (conceptualization, data curation, formal analysis, investigation, methodology, writing – review and editing), VLE (data curation, investigation, writing – review and editing), CME (data curation, writing – review and editing), LAC (conceptualization, formal analysis, funding acquisition, methodology, supervision, visualization, writing – original draft preparation).

## Abstract

Adipocytes express renin-angiotensin system (RAS) components, including angiotensinogen (*Agt*), the precursor to angiotensin II (AngII), and the angiotensin type 1a receptor (*AT1aR*). The RAS contributes to atherosclerosis, and AngII infusion causes abdominal aortic aneurysm (AAA) formation. Obesity, an established vascular risk factor, exhibits a dysregulated RAS. We studied effects of adipocyte *Agt* or *AT1aR* deficiency on diet-induced atherosclerosis and AngII-induced AAAs in male low density lipoprotein receptor (*Ldlr*) deficient mice. For atherosclerosis, male control or adipocyte *Agt/AT1aR* deficient mice were fed Western diet for 3 months. There was no effect of adipocyte *Agt* or *AT1aR* deficiency on body weight, serum cholesterol concentrations, or atherosclerotic lesions. For AngII-induced AAAs, male control and adipocyte *Agt/AT1aR* deficient *Ldlr^-/-^* mice fed Western diet were infused with AngII. Adipocyte *Agt* deficiency had no effect on body weight, serum cholesterol concentrations, abdominal aortic lumen diameter, AAA incidence, or atherosclerosis. Control, but not adipocyte *AT1aR* deficient mice lost weight during AngII infusion. The size of adipocytes in white fat was increased in adipocyte *AT1aR* deficient mice with no significant influences on abdominal aortic lumen diameter, AAA incidence, or atherosclerosis. To define mechanisms, male *Ldlr^-/-^* mice were fed standard or Western diet (1 or 3 months) and *Agt* or *AT1aR* mRNA abundance quantified in periaortic fat (PAF). *Agt* mRNA abundance in abdominal PAF increased over time in both diet groups, with modest diet-induced reductions in thoracic PAF *Agt* mRNA abundance. There was an effect of diet duration on *AT1aR* mRNA abundance in thoracic PAF, and an interaction between diet and time on abdominal PAF *AT1aR* mRNA abundance. In conclusion, adipocyte *Agt* or *AT1aR* deficiency had minimal effects on atherosclerosis or AngII-induced AAAs. However, adipocyte *AT1aR* deficient mice exhibited increased adipocyte size. Diet-induced regulation of *Agt* or *AT1aR* mRNA abundance in PAF may have contributed to these findings.

## Introduction

The renin-angiotensin system (RAS) has been implicated in cardiovascular diseases associated with obesity and dysregulated adipocyte function, including hypertension (1), atherosclerosis (2) and abdominal aortic aneurysms (AAAs) (3). Adipocytes express components of the RAS necessary to produce (angiotensinogen, *Agt*) or respond (angiotensin type 1a receptor, *AT1aR*) to angiotensin II (AngII).(1, 4) Studies suggest that adipocytes contribute approximately 25% (5) to the circulating pool of *Agt* in adult male C57BL/6 mice, and that adipose-derived *Agt* is important in the systemic production of AngII to promote obesity-hypertension (6). In addition to hypertension, *Agt* has been implicated in the development of atherosclerotic lesions (7). Both whole body *Agt* deficiency (8, 9) and liver-specific *Agt* deficiency (10) reduced atherosclerosis in hypercholesterolemic mice. However, reduced atherosclerosis in these experimental models was associated with reductions of the renal RAS, but not the systemic RAS (8, 10), suggesting that non-hepatic sources of *Agt* may directly or indirectly influence atherosclerosis.

Previous studies demonstrated that whole body deficiency of *AT1aR* profoundly reduced diet-induced atherosclerosis (2) and abolished AngII-induced AAAs (11). However, deficiency of *AT1aR* in smooth muscle (12), endothelial (12) or bone marrow cells (11) had no effect on AngII and/or diet-induced atherosclerosis or AAAs. Obesity, an inflammatory condition associated with adipocyte hypertrophy (13), is a risk factor for atherosclerosis, and previous studies demonstrated that obesity promoted AngII-induced AAAs (14). Hypertrophy of abdominal adipocytes in high fat-fed obese male mice was associated with increased markers of periaortic adipose tissue inflammation and higher infiltration of macrophages into visceral adipose tissue of obese mice that were susceptible to AngII-induced AAAs (14). We demonstrated previously that adipocyte *AT1aR* deficiency promoted hypertrophy of adipocytes from low fat (LF)-fed mice (6), an effect that would be predicted to augment atherosclerosis and AAA formation. However, while studies suggest a role for *AT1aR* in peri-vascular and visceral adipose tissue as contributors to atherosclerosis (15) and/or AAA formation (16), it is unclear if these effects were mediated by direct actions of AngII at adipocyte *AT1aR*.

The purpose of this study was to define a role for adipocyte-derived *Agt* or *AT1aR* in the development of diet-induced atherosclerosis or AngII-induced AAAs, two vascular diseases that are associated with an activated RAS (17, 18) and for which obesity is a risk factor (14, 19, 20). We hypothesize that adipocyte RAS components, specifically *Agt* and/or *AT1aR*, contribute to the development of diet-induced atherosclerosis and AngII-induced AAAs. To test this hypothesis, we used transgenic mice with genetic deletion of *Agt* or *AT1aR* in adipocytes through Cre-mediated deletion driven by fatty acid (*aP2*) binding protein. Creation of adipocyte *Agt* or *AT1aR* deficient mice using this approach was demonstrated in previous studies (5, 21) to be effective and relatively selective to adipose tissue. To induce atherosclerosis and promote AngII-induced AAAs, all studies were performed in control or adipocyte *Agt* or *AT1aR* deficient low density lipoprotein receptor (*Ldlr*) deficient mice fed Western diet for 3 months (atherosclerosis) or 5 weeks (AngII-induced AAAs). Results suggest that adipocyte *AT1aR* deficiency resulted in resistance to AngII-induced reductions in body weight and adipose tissue mass, with hypertrophied adipocytes of *AT1aR* deficient mice exhibiting resistance to body weight lowering from infusion of AngII. However, these effects were not associated with changes in measures of AngII-induced AAAs, with minimal effects of adipocyte *Agt* or *AT1aR* deficiency on diet-induced atherosclerosis.

## Methods

### Ethics Statement

All experimental procedures, including determination of humane endpoints for aneurysm studies, were reviewed and approved by the Institutional Animal Care and Use committee at the University of Kentucky.

### Animals and Diets

Previously established *Ldlr^-/-^* mice with loxP sites flanking exon 2 of the *Agt* gene were bred to transgenic male *Ldlr^-/-^* mice expressing *^aP2^*-Cre recombinase (5, 6). Details on the efficiency and selectivity of *^aP2^*-Cre deletion of *Agt* have been previously described (5). Previously established *AT1aR^fl/fl^* mice (2, 11, 12, 22, 23) on an *Ldlr^-/-^* background were bred to transgenic male mice expressing *^aP2^*-Cre recombinase (21). Details on the efficiency and selectivity of *^aP2^*-Cre deletion of *AT1aR* have been previously described (21). All studies were performed on an *Ldlr^-/-^* background to study diet-induced atherosclerosis and AngII-induced AAAs.(24, 25) Male mice (2 months of age) were fed Western diet (TD.88137, Teklad, Dublin, VA) for 3 months (for diet-induced atherosclerosis) or for one week prior to implantation of osmotic minipumps and through study duration (4 weeks for AngII-induced AAAs). For diet-induced regulation of *Agt* or *AT1aR* mRNA abundance, male *Ldlr^-/-^* mice (2 months of age; n=7 mice/diet/time point) were fed standard murine diet (Chow) or Western diet for 1 or 3 months.

### AngII-induced AAAs

At 2 months of age, mice of each genotype were anesthetized (isoflurane, to effect) for implantation of osmotic minipumps (Model 2004, DURECT Corporation, Cupertino, CA) for infusion of saline or AngII (1,000 ng/kg/min) for 28 days as previously described (25). Some mice exhibited rupture of aneurysms in the aorta during the 28 days of AngII infusion. Because of the possibility of aortic rupture, mice were monitored daily by laboratory personnel trained to recognize signs of impending aortic rupture during AngII infusions for impending signs of rupture, which included loss of lower limb movement, lethargy and loss of more than 20% body weight (measured weekly). Mice exhibiting signs of impending rupture are euthanatized within 30 minutes and then aortic rupture is defined post-mortem. However, for these studies, no mice that died prior to study endpoint (28 days of AngII infusions) exhibited impending signs of aortic rupture upon daily inspection. For studies using mice with adipocyte *Agt* deficiency, six control (*Agt^fl/f^l*) mice (out of an n = 32 mice) died of confirmed aortic rupture during AngII infusions, while five mice with adipocyte *Agt* deficiency (*Agt^Ap2^*) (out of an n = 33 mice) died of confirmed aortic rupture during AngII infusions. For studies using mice with adipocyte *AT1aR* deficiency, one control (*AT1aR^fl/fl^*) mouse (out of n=20) died of confirmed aortic rupture during AngII infusions, and three mice (out of n=24) with adipocyte AT1aR deficiency (*AT1aR^aP2^*) exhibited aortic rupture during AngII infusions.

### En face quantification of atherosclerotic lesions

The length of the aorta was removed and the intimal surface exposed by a longitudinal cut through the inner curvature of the aortic arch and down the anterior aspect of the remaining aorta (2). A cut was made through the greater curvature of the aortic arch to the subclavian branch. The tissue was pinned to a dark surface and arch areas were defined by drawing a 3-mm line from the left subclavian artery. The intimal area of the aortic arch was defined as the region from the orifice of innominate artery to the orifice of the left subclavian artery. Atherosclerotic areas (white tufts) were quantified by drawing a line around the borders and summing the total area of each lesion. Summed lesions were divided by the total arch area to calculate the percent lesion area. Measurements were performed using Nikon NIS-Elements Version 5.11.

### Measurement of Systolic Blood Pressure

Systolic blood pressure was quantified on conscious mice during week 3 of infusions using a Visitech system (BP-2000, Apex, NC) as previously described (26). Blood pressure was recorded for 5 consecutive days at the same time/day. After two days of acclimation to the system, recordings were averaged over 3 days and mice having three successful recordings were included for statistical analysis.

### In Vivo Ultrasound of Abdominal Aortic Lumen Diameter

We used ultrasound (Vevo 2100 high resolution imaging system, 55MHz probe, VisualSonics, Inc) on anesthetized (2-3% vol/vol isoflurane) mice as previously described (27). Abdominal hair was removed by shaving and applying a depilatory cream. Lumen diameter measurements were quantified by two observers blinded to the experimental design using Vevo Lab 1.7.2 software. Mice that completed the 28-day infusion protocol were included in chronic measurements of abdominal aortic lumen diameter.

### Quantification of AAA Incidence

To define mice with an AAA, cleaned aortas were pinned to a black background and two observers blinded to the experimental design assigned mice to those having AAA pathology. The percentage of mice with an aneurysm, which included mice that died from aortic rupture confirmed post-mortem, was calculated as AAA incidence for each group.

### Plasma/Serum Measurements

At study endpoint, blood was obtained by cardiac puncture on anesthetized mice for plasma (EDTA 5 mM; aprotinin 500 KIU/ml) to quantify renin concentrations, or for serum (BD Microtainer SST tubes) to quantify total serum cholesterol or triglyceride concentrations. Plasma was obtained following centrifugation (5,000 rpm, 10 min, 4°C) and stored at −80°C. Sera was obtained following centrifugation (13,000 rpm, 4 min, 37°C) and stored at −80°C. Plasma renin concentrations were quantified as previously described (27). Mouse plasma (8 µl) was incubated in buffer (100 mM Tris/HCl, pH 8.5, 20 mM EDTA, 1 mM PMSF) with and without the renin substrate *Agt* (purified from nephrectomized rat plasma) for 30 minutes (37°C). Angiotensin I (AngI) in plasma was quantified using an ELISA (IBL, #BI59131; sensitivity of 0.14 ng/ml). Background levels of AngI (without addition of Agt) were subtracted from levels when incubated with substrate to obtain plasma renin concentrations.

### Quantification of Gene Expression

To define influences of study duration or diet on *Agt* or *AT1aR* mRNA abundance in periaortic fat (PAF), male *Ldlr^-/-^* mice were fed standard murine diet (Chow) or Western diet for 1 or 3 months. At study endpoint, mice were euthanatized for tissue (PAF) harvest. RNA was extracted using a Maxwell Rapid Sample Concentrator (RSC) simply RNA Tissue kit (REF#AS1340, Promega, Madison, WI). RNA was reverse transcribed to cDNA using Supermix (CAT#95048-500, Quanta Biosciences, Gaithersburg, MD). cDNA was diluted (1:10) and mRNA abundance was quantified by RT-PCR using SYBER Green FastMix (CAT#95071-012, Gaithersburg, MD) on a BioRad quantitative real-time PCR thermocycler (CF96 Real-Time system, BioRad, Hercules, CA). mRNA abundance was determined using the ΔΔCt method and normalized to 18S RNA as a housekeeping gene. Primer sequences are illustrated as follows:

Primer sequences for RT-PCR quantification of *Agt* or *AT1aR* mRNA abundance.

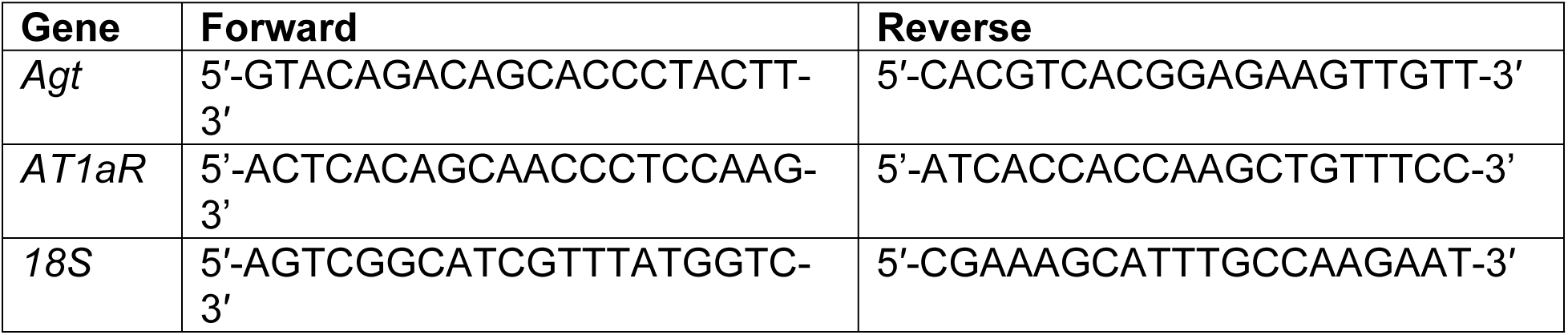

### Quantification of Adipocyte Size and Number

Sections of formalin (10% w/v) fixed pieces of retroperitoneal tissue were stained with hematoxylin and eosin (H&E) (21). Images of slides were obtained at 10x magnification and using the “detect edges”, image threshold, and object count features of NIS Elements software (Nikon, Instruments, Inc., Tokyo, Japan), the area of each adipocyte within a 700 x 700 µm measurement frame was quantified. Three measurement frames on each slide and 3 slides per genotype group were analyzed for morphology.

### Statistical Analyses

For illustration, data are presented as mean ± standard error of the mean (SEM). Data sets were tested for normality (Shapiro-Wilk) and equal variance (Brown-Forsythe). To define differences between groups (normally distributed data), data were analyzed using unpaired Student *t*-test for two sample comparisons, by repeated measures two-way ANOVA (ultrasound, body weights), or by two-way ANOVA (genotype, treatment) for multiple group comparisons using Holm-Sidak post-hoc tests. For categorical data (AAA incidence), data were analyzed using Fisher’s Exact test. Statistical significance was defined as P<0.05. Statistical analyses were performed using GraphPad Prism 8.

## Results

### Neither adipocyte Agt nor AT1aR deficiency regulate diet-induced atherosclerosis

Control (*Agt^fl/fl^*) and adipocyte *Agt* deficient (*Agt^aP2^*) male mice were fed Western diet for 3 months to induce atherosclerosis. There was no difference in body weight (Fig. 1A), fat mass (Table 1), serum cholesterol (Fig. 1B) or triglyceride concentrations (Table 1) between *Agt^fl/fl^* control or *Agt^aP2^* adipocyte deficient mice. Plasma renin concentrations were quantified because previous studies demonstrated that adipocytes contributed to the systemic *Agt* pool and could thereby influence systemic renin concentrations.(5, 6) Plasma renin concentrations were increased in *Agt^aP2^* mice compared to *Agt^fl/fl^* controls, but there were no significant differences between groups (Table 1; p = 0.063). Atherosclerotic lesion surface area in the aortic arch was not significantly different between genotypes (Fig. 1C, p = 0.529).

**Figure 1.**
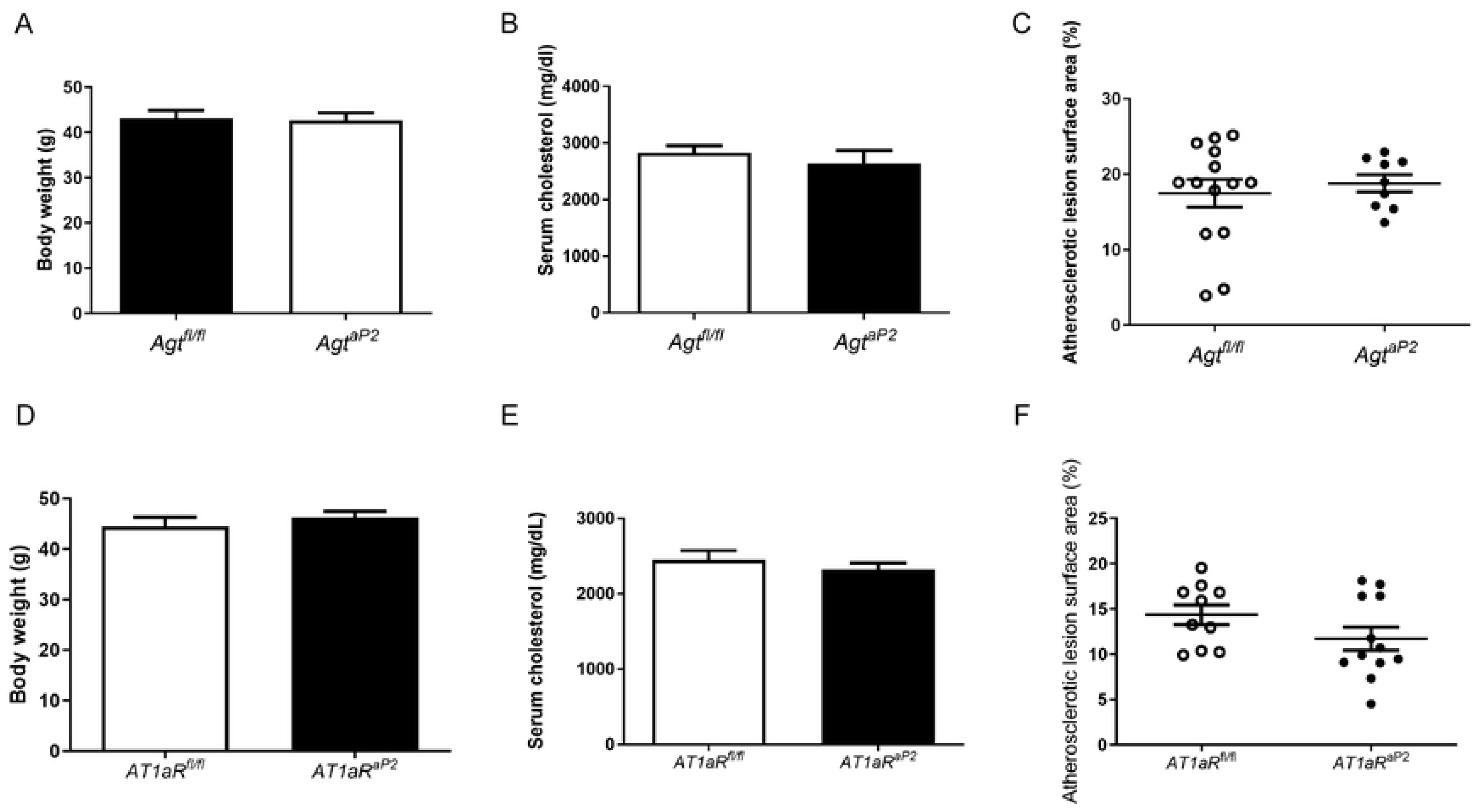
Effects of adipocyte *Agt* or *AT1aR* deficiency on diet-induced atherosclerosis of *Ldlr^-/-^* male mice. Male control (*Agt^fl/fl^*, *AT1aR^fl/fl^*) or adipocyte-deficient (*Agt^aP2^*, *AT1aR^aP2^*) *Ldlr^-/-^* mice were fed Western diet for 3 months. A-C, control (n = 14) and adipocyte *Agt* deficient mice (n=9). A, Body weight of control and adipocyte *Agt* deficient mice. B, Serum cholesterol concentrations. C, Atherosclerotic lesion percent surface area of the aortic arch. D-E, control (n=19) and adipocyte *AT1aR* deficient mice (n=21). D, Body weight. E, Serum cholesterol concentrations. F, Atherosclerotic lesion percent surface area of the aortic arch. Data are mean ± SEM.

**Table 1.**
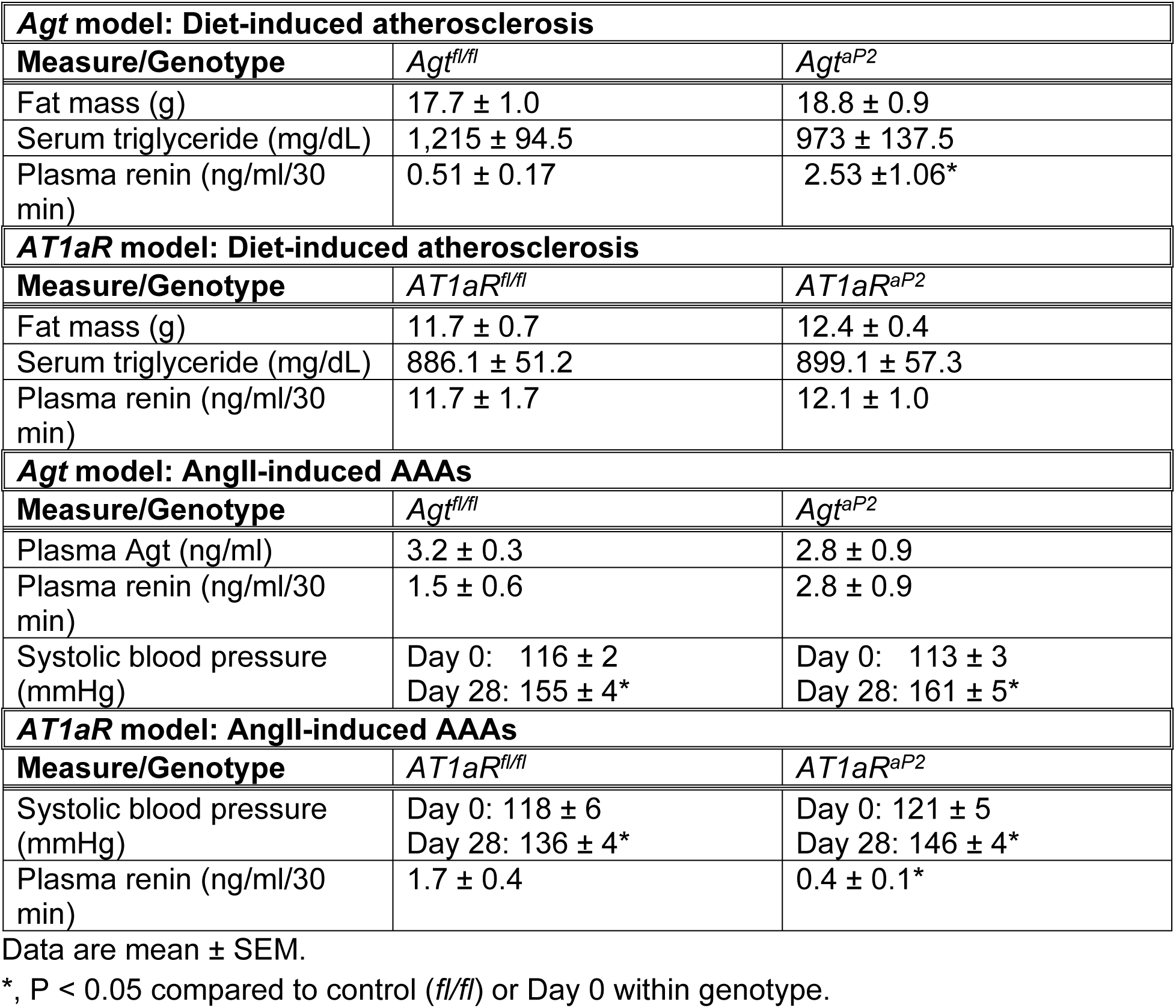
Characteristics of mice with adipocyte deficiency of *Agt* or *AT1aR*.

Similarly, control *AT1aR^fl/fl^* and adipocyte *AT1aR* deficient (*AT1aR^aP2^*) male mice were fed Western diet for 3 months. There was no significant difference in body weight (Fig. 1D; p = 0.397), fat mass (Table 1), serum cholesterol (Fig. 1E; p = 0.385) or triglyceride concentrations (Table 1; p = 0.872) between *AT1aR^fl/fl^* control or adipocyte *AT1aR^aP2^* deficient mice. As *AT1aR* mediate negative feedback of AngII on kidney renin secretion (28, 29), we quantified plasma renin concentrations which were not significantly different between genotypes (Table 1). Atherosclerotic lesion surface area in the aortic arch was not significantly different between genotypes (Fig. 1F, p = 0.142).

### Neither Adipocyte Agt nor AT1aR deficiency regulate AngII-induced AAAs

To study the influence of adipocyte *Agt* on AngII-induced AAAs, adipocyte *Agt^aP2^* deficient and *Agt^fl/fl^* control *Ldlr^-/-^* mice fed Western diet were infused with AngII for one month. Body weight at study endpoint was not significantly different between genotypes (Fig. 2A; p = 0.163). Plasma concentrations of *Agt* (Table 1; p = 0.346), renin (Table 1; p = 0.304) and serum cholesterol concentrations (Fig. 2B; p = 0.320) were not significantly different between genotypes. Systolic blood pressures of AngII-infused mice were increased compared to day 0 in both genotypes (p< 0.0001), with no significant differences between genotypes (Table 1; p = 0.956). Abdominal aortic lumen diameters of AngII-infused mice were significantly increased compared to day 0 (p < 0.001) in both genotypes with no significant differences between genotypes (Fig. 2C; p = 0.806). AAA incidence was similar between genotypes (Fig. 2D; p = 0.602), as was the percent of the aortic arch covered by atherosclerotic lesions (Fig. 2E; p = 0.988).

**Figure 2.**
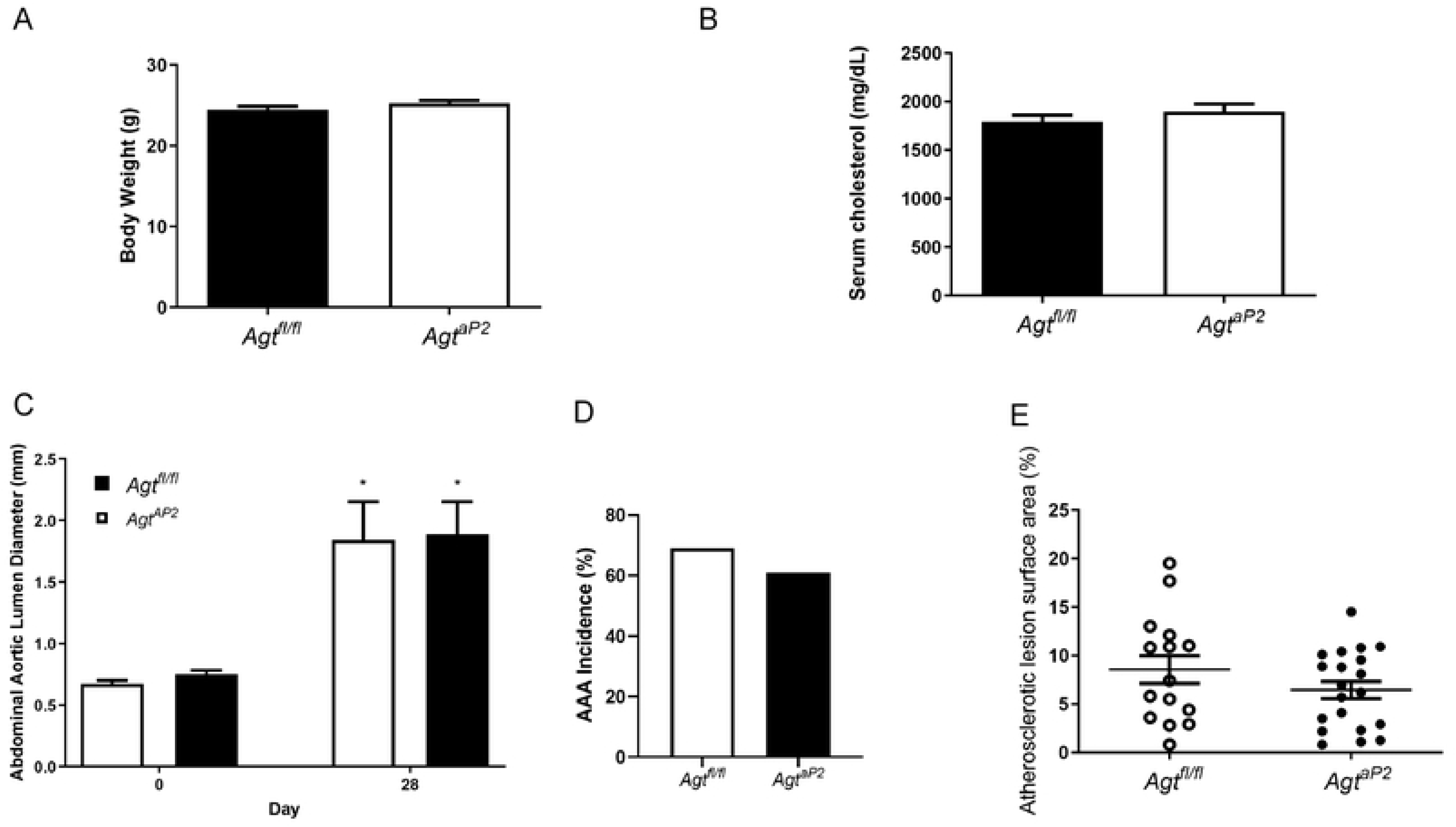
Effects of adipocyte *Agt* deficiency on AngII-induced AAAs. Male control (*Agt^fl/fl^*; n=32) or adipocyte deficient (*Agt^aP2^*; n=33) mice were fed Western diet for one week before implantation of osmotic minipumps containing AngII (1,000 ng/kg/min) for 28 days. A, Body weight. B, Serum cholesterol concentrations. C, Abdominal aortic lumen diameters of control (n=25) and adipocyte *Agt* deficient mice (n=26) on day 0 and 28 of AngII infusions. D, AAA incidence of AngII-infused control and adipocyte *Agt* deficient mice. E, Atherosclerotic lesion surface area of the aortic arch of control and adipocyte *Agt* deficient AngII-infused mice. Data are mean ± SEM. *, P < 0.05 compared to day 0 within genotype.

To study effects of adipocyte *AT1aR* deficiency on AngII-induced AAAs, we included groups of *AT1aR^fl/fl^* and *AT1aR^aP2^* mice infused with saline or AngII, as previous studies demonstrated alterations in adipocyte size in non-infused adipocyte *AT1aR* deficient mice (21). Body weight increased significantly (p < 0.0001) in both genotypes, with no significant differences between genotypes (Fig. 3A; p = 0.133).

**Figure 3.**
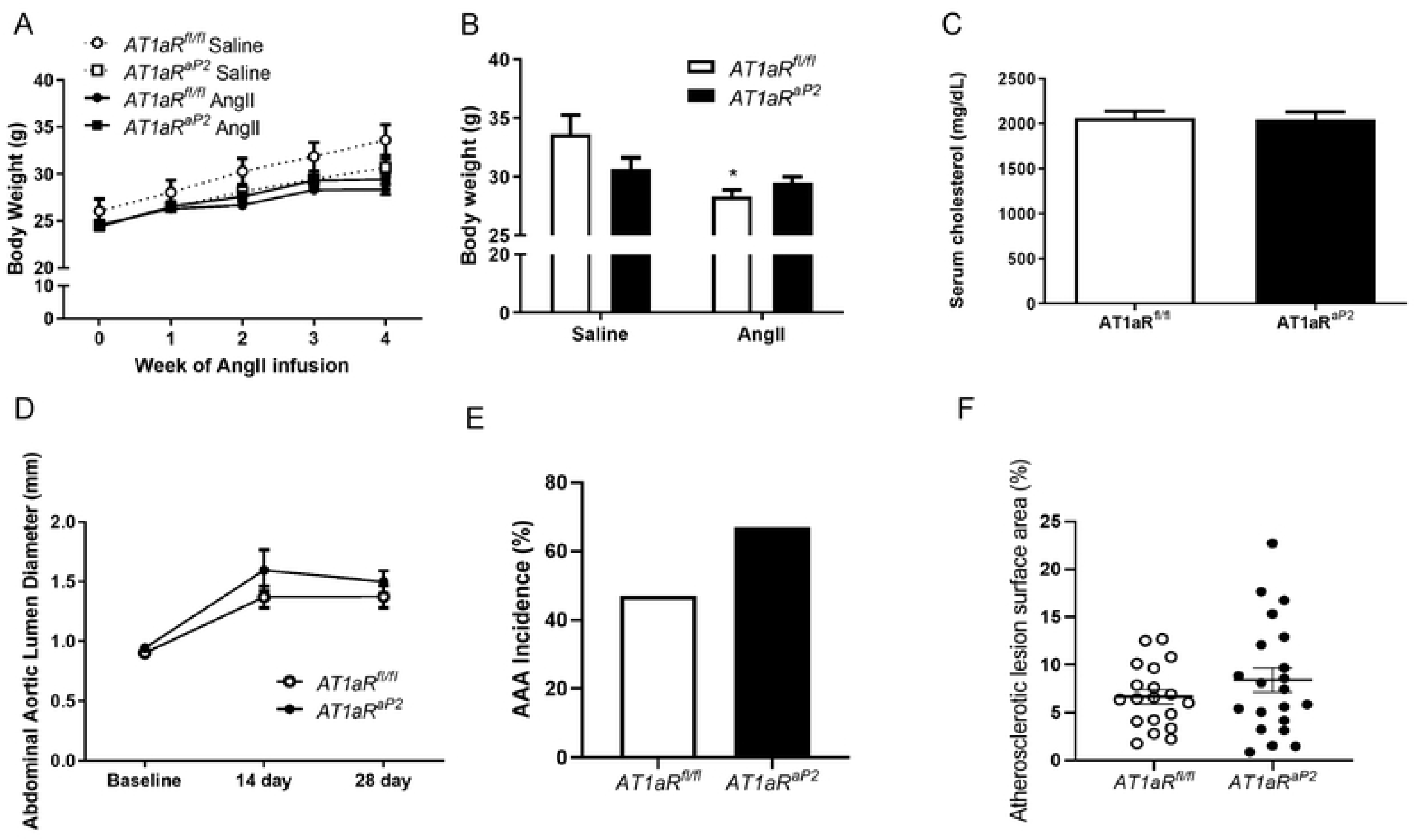
Effects of adipocyte *AT1aR* deficiency on AngII-induced AAAs. Male control (*AT1aR^fl/fl^*; n=20) or adipocyte deficient (*AT1aR^aP2^*; n=24) mice were fed Western diet for one week before implantation of osmotic minipumps containing saline or AngII (1,000 ng/kg/min) for 28 days. A, Body weight of mice of each treatment and genotype over duration of infusions. B, Body weight of mice from each group at study endpoint. C, Serum cholesterol concentrations of AngII-infused mice of each genotype. D, Abdominal aortic lumen diameters of AngII-infused mice of each genotype at baseline, 14 and 28 days of AngII infusion. E. AAA incidence of AngII-infused mice of each genotype (%). F, Atherosclerotic lesion surface area of the aortic arch of AngII-infused mice of each genotype. Data are mean ± SEM.

However, there was a significant interaction between duration of infusion and genotype (p = 0.022). We analyzed body weight at study endpoint between treatments (saline, AngII) and genotypes (*AT1aR^fl/fl^*, *AT1aR^aP2^*; Fig. 3B). On day 28 of infusions, body weight was modestly, but not significantly (p = 0.06) decreased in saline *AT1aR^aP2^* mice compared to saline *AT1aR^fl/fl^* mice (Fig. 3B). Moreover, body weight was decreased significantly (p < 0.001) by infusion of AngII for 28 days in *AT1aR^fl/fl^*, but not AngII-infused *AT1aR^aP2^* mice (Fig. 3B), with a significant interaction (p = 0.023) between genotypes. Infusion of AngII resulted in a significant increase in systolic blood pressures of both genotypes (Table 1; p < 0.001), with no differences between genotypes (p = 0.223). Plasma renin concentrations were significantly decreased in AngII-infused *AT1aR^aP2^* mice compared to AngII-infused *AT1aR^fl/fl^* controls (Table 1; p = 0.003).

However, serum total cholesterol concentrations were not significantly different between AngII-infused *AT1aR^fl/fl^* and *AT1aR^aP2^* mice (Fig. 3C; p = 0.888). Abdominal aortic lumen diameter increased significantly in both genotypes of AngII-infused mice over time (Fig. 3D; p < 0.0001), with no significant differences between genotypes (p = 0.182). AAA incidence increased modestly (from 47 to 67%), but not significantly (p = 0.324) in AngII-infused *AT1aR^aP2^* mice compared to *AT1aR^fl/fl^* controls (Fig. 3E). The percent of the aortic arch covered by atherosclerotic lesions was not significantly different between AngII-infused *AT1aR^fl/fl^* and *AT1aR^aP2^* mice (Fig. 3F; p = 0.271).

Because previous studies indicated a novel phenotype of adipocyte hypertrophy in LF-fed *AT1aR^aP2^* mice (21) and based on results suggesting resistance of *AT1aR^aP2^* mice to reductions in body weight from AngII infusion (Fig. 3A,B), we analyzed adipocyte size in retroperitoneal white adipose tissue of saline and AngII-infused *AT1aR^fl/fl^* and *AT1aR^aP2^* mice (Fig. 4A-D). Histograms of adipocyte size (µm^2^) *versus* cell number for each treatment group illustrated a shift of adipocytes to larger sizes in saline and AngII-infused *AT1aR^aP2^* mice compared to *AT1aR^fl/fl^* controls (Fig. 4), with most pronounced increases of adipocyte size of AngII-infused *AT1aR^aP2^* mice compared to AngII-infused *AT1aR^fl/fl^* controls (Fig. 4B). Moreover, despite no differences in end-point body weights between genotypes of AngII-infused mice (Fig. 3B), the weight of retroperitoneal white adipose tissue was decreased significantly in AngII-infused *AT1aR^fl/fl^* mice compared to saline, but not in AngII-infused *AT1aR^aP2^* mice compared to saline (Fig. 4D; p = 0.040). Thus, increased adipocyte size was observed in retroperitoneal adipose tissue exhibiting reduced mass of AngII-infused *AT1aR^aP2^* mice compared to *AT1aR^fl/fl^* controls.

**Figure 4.**
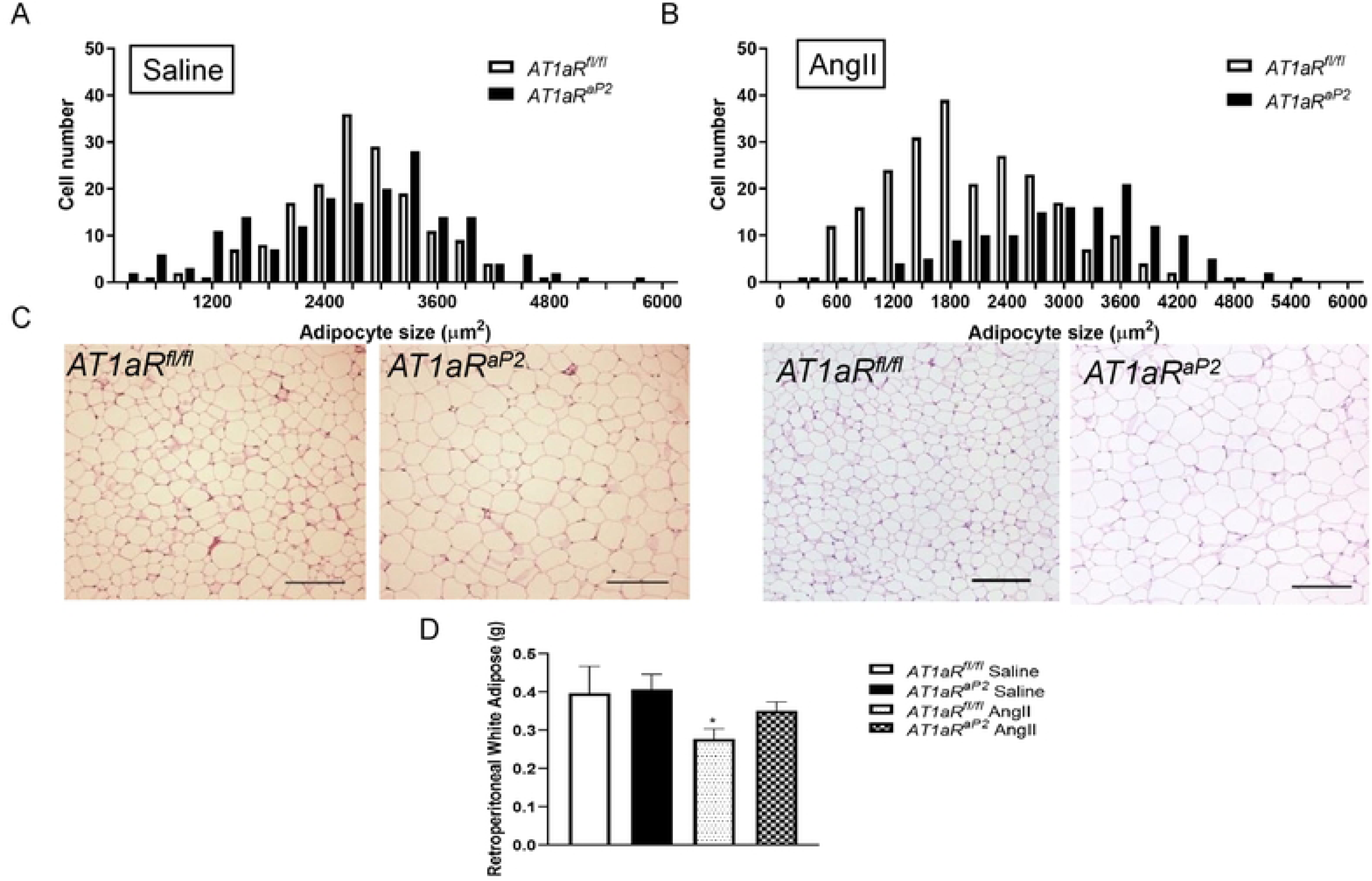
Effects of adipocyte *AT1aR* deficiency on adipocyte size in white adipose tissue. Male control (*AT1aR^fl/fl^*) or adipocyte deficient (*AT1aR^aP2^*) mice were fed Western diet for one week before implantation of osmotic minipumps containing saline or AngII (1,000 ng/kg/min) for 28 days. A, Cell number *versus* adipocyte size in retroperitoneal white adipose tissue sections from saline-infused mice (n=9) of each genotype. B, Cell number *versus* adipocyte size in retroperitoneal white adipose tissue sections of AngII-infused mice of each genotype. C, Representative images of retroperitoneal white adipose tissue sections stained with H&E from saline (left) and AngII (right)-infused mice (n=9) of each genotype. D, Weight (g) of retroperitoneal white adipose tissue of saline and AngII-infused mice of each genotype. A,B, Data represent three measurement frames on each slide and three slides/genotype. D, Data are mean ± SEM. *, P < 0.05 compared to saline within genotype.

### Influences of time and diet on Agt or AT1aR mRNA abundance in thoracic and abdominal PAF

Previous studies demonstrated age-or diet-induced regulation of adipose *Agt* mRNA abundance in adipose tissue (30–32), or a potential role of *AT1aR* in the regulation of adiposity in response to a high fat diet (33). Therefore, we investigated influences of study time point duration or Western diet feeding on *Agt* or *AT1aR* mRNA abundance in thoracic and abdominal PAF. We chose PAF as an adipose source since this tissue has been previously suggested to influence atherosclerosis (34, 35) and/or AAA formation (14, 36–38). Moreover, there is no anatomic barrier between PAF and the aortic wall where both atherosclerotic lesions and AAAs form, and PAF expresses both of these RAS components (39, 40). Male *Ldlr^-/-^* mice (2 months of age) were fed Chow or Western diet for 1 or 3 months, representing durations of study designs used for AngII-induced AAAs (1 month of diet or 3 months of age) or diet-induced atherosclerosis (3 months of diet or 5 months of age). At both time points, *Agt* mRNA abundance was modestly, but not significantly (p = 0.110) reduced in thoracic PAF from mice fed Western diet compared to Chow (Fig. 5A). In contrast, *Agt* mRNA abundance was increased significantly in abdominal PAF from 3 month compared to 1 month mice of each diet group (Fig. 5B, p = 0.047), with no differences between diet groups at either time point (p = 0.579). *AT1aR* mRNA abundance decreased significantly (p = 0.001) in thoracic PAF from mice fed Western diet for 3 months compared to 1 month, with no significant differences in thoracic PAF *AT1aR* mRNA abundance between 1 and 3 months of Chow feeding (Fig. 5C, p = 0.164). In abdominal PAF, there was a significant interaction (p = 0.007) between diet and time for *AT1aR* mRNA abundance (Fig. 5D), with elevations of abdominal PAF *AT1aR* mRNA abundance over time in mice fed Chow, *versus* reductions of abdominal PAF *AT1aR* mRNA abundance in mice fed Western diet for 3 compared to 1 month.

**Figure 5.**
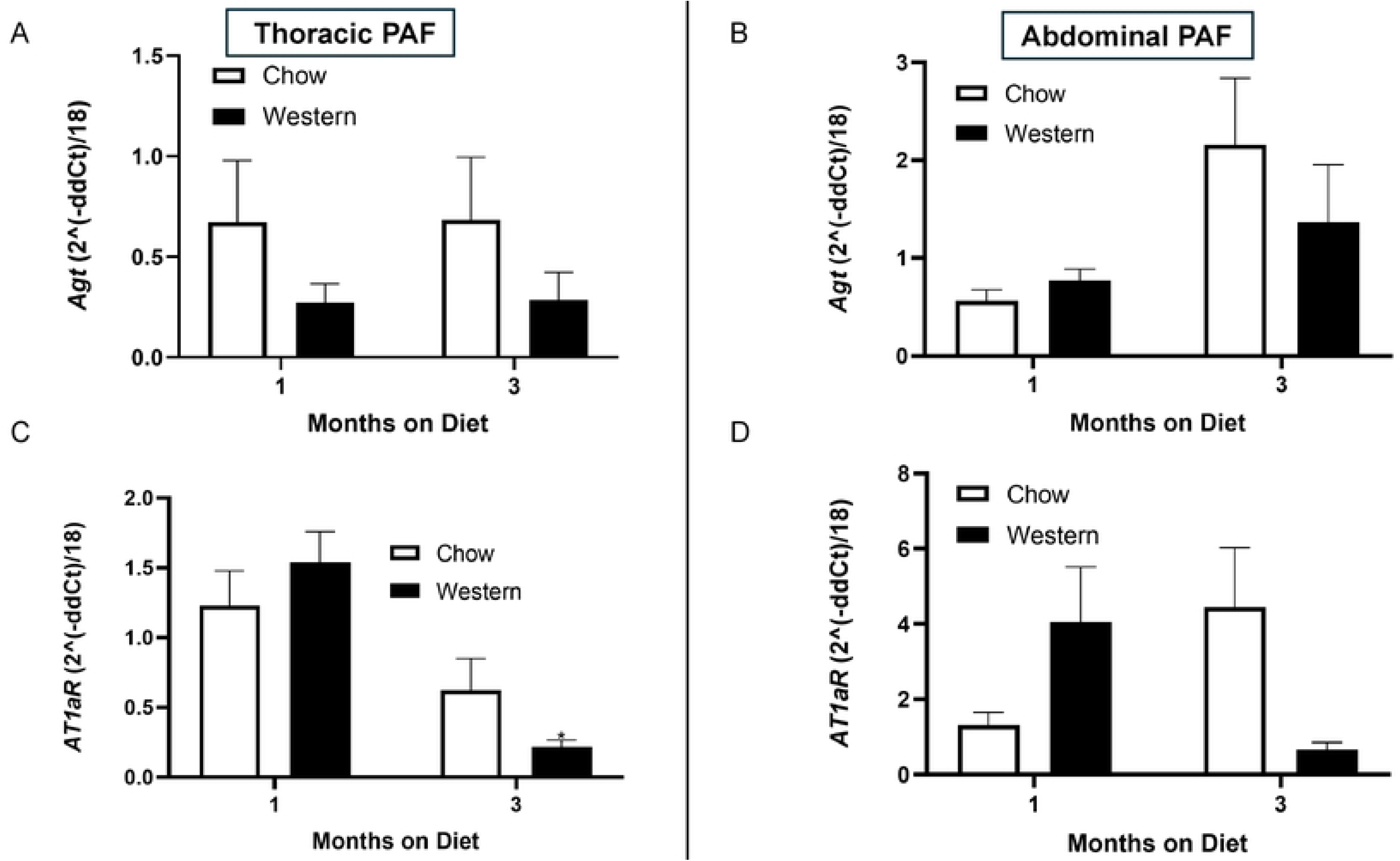
Effects of diet and/or time on *Agt* or *AT1aR* mRNA abundance in thoracic and abdominal periaortic fat (PAF). Male *Ldlr^-/-^* mice were fed standard murine diet (chow) or Western diet for 1 or 3 months. A, *Agt* mRNA abundance in thoracic PAF. B, *Agt* mRNA abundance in abdominal PAF. C, *AT1aR* mRNA abundance in thoracic PAF. D. *AT1aR* mRNA abundance in abdominal PAF. Data are mean ± SEM from n=7 mice fed each diet and at each time point. *, P < 0.05 compared to Chow at same time point.

## Discussion

We found that adipocyte-specific deficiency of RAS components, namely *Agt* or *AT1aR*, had no effect on diet-induced atherosclerosis in *Ldlr^-/-^* mice. Using a different vascular disease model induced by infusion of AngII to hypercholesterolemic male mice, deficiency of *Agt* in adipocytes had no influence on AAA formation. Moreover, although adipocyte *AT1aR* deficiency modestly increased measures of AAA formation and atherosclerosis, these differences were not statistically significant. Notably, adipocyte *AT1aR* deficiency resulted in an interesting phenotype of adipocyte hypertrophy despite reductions in mass of a white adipose depot in AngII-infused mice. Additionally, these mice resisted the body weight loss typically induced by infusion of AngII in Western diet-fed *Ldlr^-/-^* mice. To understand mechanisms influencing the role of *Agt* or *AT1aR* in adipocytes on vascular diseases, we quantified mRNA abundance in PAF over the study durations and in response to Western diet. Results suggest that Western diet-induced reductions of *AT1aR* mRNA abundance may have contributed to a lack of a significant effect of adipocyte *AT1aR* deficiency on the development of atherosclerosis or AAAs.

Previous studies demonstrated efficient *^aP2^*^-^Cre-driven deletion of *Agt* in white and brown adipose tissue of male C57BL/6 mice, with minimal influences on *Agt* mRNA abundance in non-adipose tissues (5). Consistent with these findings, our data show that AngII-infused adipocyte *Agt* deficient mice exhibit modest reductions in plasma *Agt* levels, similar to reductions observed in low fat-fed male *Agt* adipocyte-deficient C57BL/6 mice compared to controls (5). Moreover, we observed significantly elevated plasma renin concentrations in adipocyte *Agt*-deficient male mice fed a Western diet for 3 months, suggesting that lower plasma AngII levels (due to reduced adipocyte-derived *Agt*) might relieve the negative feedback on renin production (41). Previous studies demonstrated that reductions in hepatic *Agt* (10) significantly decreased atherosclerosis in *Ldlr^-/-^* male mice. These results demonstrated that endogenous (hepatic) *Agt* contributes to the development of experimental diet-induced atherosclerosis. However, in the present study, despite modest changes in the systemic RAS of mice with adipocyte *Agt* deficiency, there were no significant differences in serum lipids or atherosclerotic lesion areas between genotypes. Thus, in contrast to hepatic-derived *Agt* (10), these results do not support adipocyte-derived *Agt* as a primary contributor to the development of diet-induced atherosclerosis.

Previous studies demonstrated that whole body deficiency of *AT1aR* markedly decreased the development of diet-induced atherosclerosis of *Ldlr^-/-^* mice (2). However, neither deficiency of endothelial (12), nor smooth muscle cell (12) *AT1aR* influenced atherosclerosis of male *Ldlr^-/-^* mice. Moreover, studies using bone marrow transplantation demonstrated that while bone marrow *AT1aR* deficiency had no effect on lesion formation, transplantation of *AT1aR^+/+^* bone marrow to *AT1aR^-/-^* recipients reduced diet-induced atherosclerosis of male *Ldlr^-/-^* mice (11). Thus, neither endothelial, smooth muscle nor bone marrow cells expressing *AT1aR* contribute to diet-induced atherosclerosis, and therefore the identity of cells expressing *AT1aR* in developing atherosclerotic lesions has not been defined. Since obesity is a risk factor for atherosclerosis and adipocytes are known to express several components of the RAS including *AT1aR* (21) instrumental in atherosclerotic lesion formation (4, 42), we examined the role of adipocyte *AT1aR* on diet-induced atherosclerosis. Using an *^aP2^*-driven Cre system, we targeted *AT1aR* deletion in adipocytes, as described previously (21). Similar to results obtained with adipocyte *Agt* deficiency, *AT1aR* deficiency in adipocytes had no significant effects on serum cholesterol concentrations or atherosclerotic lesion formation. Thus, in addition to previously examined cell types, adipocyte *AT1aR* are not primary mediators of diet-induced atherosclerosis.

Previous studies demonstrated that infusion of AngII to *Ldlr^-/-^* mice increased mRNA abundance of *Agt* and *AT1aR* in adipose tissue (43). These results suggested a potential role for adipocyte *Agt* and/or *AT1aR* in the formation of AAAs from AngII infusion to hypercholesterolemic *Ldlr^-/-^* mice. Several studies have invoked a potential role for abdominal periaortic fat in the development of human and experimental AAAs.(14, 37, 38, 44, 45) Notably, periaortic fat is a potential source of *Agt* (39) and also expresses *AT1aR* (15, 16, 40, 46). However, our results do not support a role for adipocyte-derived *Agt* in AngII-induced AAAs.

We demonstrated previously that obesity promotes the formation of AngII-induced AAAs associated with inflammation of abdominal periaortic fat (14), supporting a role for adipose tissue in AAA development. Previous results demonstrated that deficiency of *AT1aR* in adipocytes resulted in a phenotype of adipocyte hypertrophy of low fat-fed mice (21). Further studies using stromal vascular cells from adipocyte *AT1aR* deficient mice demonstrated impaired differentiation of preadipocytes to mature adipocytes, while conversely exposure of differentiating preadipocytes to AngII increased Oil Red O staining of mature adipocytes (21). Similar effects of AngII to promote adipocyte differentiation were reported by others (47–49). Taken together, these results suggest that AngII promotes formation of new adipocytes through *AT1aR*.

In the present study white adipose tissue of adipocyte *AT1aR* deficient mice exhibited hypertrophy. As observed in previous studies (21), reductions of adipocyte differentiation in response to AngII infusions of adipocyte *AT1aR* deficient mice may have contributed to hypertrophy of remaining adipocytes. Since obesity is associated with hypertrophied, inflamed adipocytes (50), and obesity has been demonstrated to promote AngII-induced AAAs (14), these results suggest that modest elevations in measures of AAA formation of adipocyte *AT1aR* deficient mice may have resulted from the influences of hypertrophied adipocytes.

A potential contributor to the lack of effect of adipocyte *Agt* or *AT1aR* on atherosclerosis and/or AAA relates to effects of Western diet to regulate expression of these genes in adipose tissue. In the present study, *AT1aR* mRNA abundance in PAF was reduced in mice fed Western diet for 3 months compared to mice fed standard murine diet (Chow). Diet-induced reductions in PAF *AT1aR* mRNA abundance may have minimized effects from adipocyte *AT1aR* deficiency on atherosclerosis and/or AAA formation. However, as there was minimal diet-induced regulation of PAF *Agt* mRNA abundance, collectively, these results do not support a role for adipocyte-derived *Agt* in vascular disease development.

In conclusion, results from this study do not support a prominent role for adipocyte-derived *Agt* or *AT1aR* in atherosclerosis or AAA formation. Thus, sources of *Agt* other than adipocytes or non-adipocyte cell types that respond to AngII through *AT1aR* mediate RAS-induced potentiation of atherosclerosis and formation of AAAs.

